# TulsiPIN: an interologous protein interactome of *Ocimum tenuiflorum*

**DOI:** 10.1101/680025

**Authors:** Vikram Singh, Gagandeep Singh, Vikram Singh

**Affiliations:** Centre for Computational Biology and Bioinformatics, Central University of Himahcal Pradesh, Dharamshala, India

**Keywords:** Interologue, interactome, *Ocimum tenuiflorium*, PPI network, colocalization, functional homogeneity

## Abstract

*Ocimum tenuiflorum*, commonly known as holy basil or tulsi, is globally recognized for its multitude of medicinal properties. However, a comprehensive study revealing the complex interplay among its constituent proteins at subcellular level is still lacking. To bridge this gap, a genome scale interologous protein-protein interaction (PPI) network, TulsiPIN, is developed using 49 template plants. The reported network consists of 13, 660 nodes and 327, 409 binary interactions. A high confidence PPI network consisting of 7, 719 nodes having 95, 532 interactions was inferred using domain-domain interaction information along with interolog based statistics, and its reliability was further assessed using functional homogeneity and protein colocalization. 1, 625 vital proteins are predicted by statistically evaluating this high confidence TulsiPIN with two ensembles of corresponding random networks, each consisting of 10, 000 realizations of Erdős-Rényi and Barabási-Albert models. Topological features of TulsiPIN including small-world, scale-free and modular architecture are inspected and found to resemble with other plant PPI networks. Finally, numerous regulatory proteins like transcription factors, transcription regulators and protein kinases are profiled in TulsiPIN and a sub-network of proteins participating in 10 secondary metabolite biosynthetic pathways is studied. We believe, the methodology developed and insights imparted would be useful in understanding regulatory mechanisms in various plant species.

## Introduction

*Ocimum tenuiflorum*, belonging to family Lamiaceae, is an aromatic shrub widely recognized for its vast array of health benefits^1^. It is commonly known as the holy basil or tulsi and has been referred as “mother medicine of nature” as well as “queen of plants” in Ayurveda, the traditional medicinal system practiced in India. The holy basil is not only regarded as a spiritual herb in Indian folklore but also worshiped as a goddess across the country. Genus *Ocimum* is highly diverse and includes more than 160 species distributed in different tropical and subtropical regions of Asia, Africa and America. So far nine species of tulsi are reported in India, among them six are native and three are exotic^2^. Tulsi has been used in clinical practice since ancient times either independently or along with several other herbs in various formulations to cure a variety of diseases or disorders^3^. Different parts of this plant including roots, stem, bark, leaves, flowers and seeds produce a number of secondary metabolites in response to different biotic and abiotic stimuli. These metabolites possess significant pharmaceutical properties and are highly valuable in healthcare, cosmetics and perfumery industries^4,5^. *Ocimum* leaves are reported to have anthelmintic, expectorant, diaphoretic and stimulating effects^6^. Its infusion is used to treat arthritis, ringworm infections, piles etc. and root decoction is used to cure malaria and urino-genital disorders^7^. Tulsi plant has also been found to be endowed with several imperative medicinal properties like, memory enhancement, neuro-protective, anti-convulsant, anti-stress, anti-spasmodic, analgesic, anti-cancer, anti-diabetic, anti-ulcer, anti-oxidant, anti-inflammatory, antimicrobial ^8^. Along with medicinal values the plant is also used as an ornament and condiment^1^.

Recently, two genomes of *Ocimum tenuiflorum* have been reported^9,10^ that provide a draft catalogue of genes and proteins present in this species. Also, transcriptome wide protein expression based studies are being carried out^9,11^ towards gaining the important insights about secondary metabolite biosynthetic pathways^12^. However, a systems level study explaining complex web of interactions among various proteins that can provide first-hand intuition about cooperation at molecular level is still lacking in tulsi. Biological systems are inherently complex and stochastic – a characteristics that can be attributed to the diverse classes of constituent small molecules like genes, proteins, metabolites etc.^13^. Interactions (molecular regulation) among these individual elements give rise to an array of emergent properties at cellular and higher levels. These emergent behaviors can not be explained solely on the basis of reductionism rather demand an exhaustive consideration of molecular interactions characterizing the entire system^14,15^. To develop a comprehensive understanding of the molecular mechanisms inherent to different biological systems, we need to investigate the underlying interactomes as a whole^16,17^. This interactomic approach has become a useful tool in identifying small functional circuits (modules) as well as key molecules characterizing complex network behavior and has enabled us to decipher the molecular mechanisms underlying biological systems^18^.

Although systems level protein-protein interaction studies are desirable, however, experimental identification of interactions is enormously expensive in terms of time, labor and cost^19^. An alternative to this problem is to computationally infer interactions between two proteins based on available protein interaction information of other species^20^. This method, called interolog approach, is based on the principle of homology and assumes that proteins encoded by orthologous genes alongside maintaining their functions, to a large extent preserve their interacting partners as well. So if a link exists between two orthologous proteins in template network, a link between corresponding proteins in the query network is also placed. Owing to quick, easy and cost effective nature of interolog approach a large number of PPI networks for important plant species including, *Arabidopsis thaliana* ^21^, *Oryza sativa* ^22^, *Zea mays* ^23^, *Manihot esculenta* ^24^, *Camellia sinensis* ^25^ etc. have been reported recently.

*O. tenuiflorum* is a highly valuable herb both economically as well as medicinally due to uncountable number of health benefits it offers. Thus, understanding the complex molecular mechanisms underlying growth, development, defense etc. of this plant will enable us in improving the yield of its essential oils as well as in producing more resistant varieties against biotic and abiotic perturbations. Towards that goal, in the present study we have constructed an interactome map of tulsi by transferring conserved interactions among proteins from a large number of template plants (see Figure 1). Each interaction in TulsiPIN is assigned a confidence score by combining the outcomes of interologous and domain-domain interaction approaches to extract a high confidence TulsiPIN. We further identified key proteins by combining global network metrics and explored the modular architecture of the predicted PPI network. To reveal the interconnections between proteins involved in 10 metabolic biosynthesis pathways, analysis of metabolic subnetwork was also preformed.

**Figure 1:**
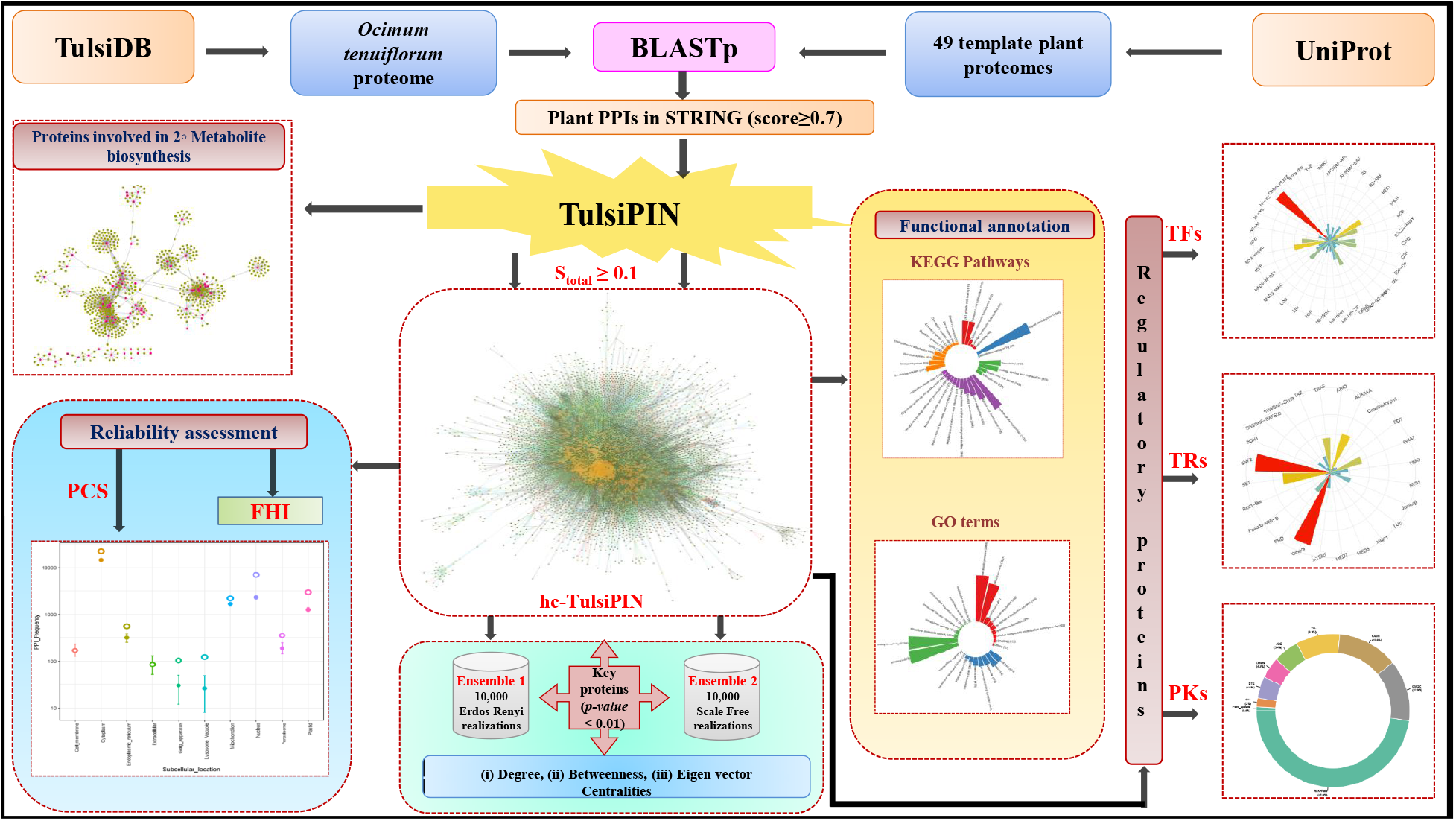
Overall workflow chart of present study

## Methods

### Data acquisition and interologous TulsiPIN construction

Protein sequence data for *O. tenuiflorum* was obtained from TulsiDb (http://caps.ncbs.res.in/Ote/) on January 2019. Protein interactions of all the 49 plant species available in STRING (11.0) database were collected^**?**^. Additionally, proteomes of these 49 plants were obtained from UniProt database^26^. Potential orthologs of tulsi proteins in individual template plant were identified by querying the tulsi proteome against the selected plant proteome using BLASTp available at NCBI^27,28^. Sequences which met all the following three conditions were considered potential orthologs: i) sequence identity *≥* 60%, ii) query coverage *≥* 80% and iii) e-value < *e*^−10^ ^24^. Then, an interologous PPI network of *O. tenuiflorum* (TulsiPIN) was constructed by mapping interactions from original template plants to corresponding orthologous proteins in tulsi. A protein pair of *O. tenuiflorum* was considered to be interacting, if both their orthologous proteins were found to be interacting in at least one of 49 template plants. The final constructed TulsiPIN was visualized using Cytoscape v3.7.0^29^.

### Domain-domain interaction prediction

Most of the proteins are estimated to be composed of multiple domains^30^ and PPIs are outcome of interaction between their constituent domain pairs^31^. Consequently, protein interactions are strongly influenced by interactions between their underlying domains and thus domain-domain interaction information can provide significant confidence to predicted protein-protein interactions^32^. For that, the domain information of *O. tenuiflorum* proteins was derived from Pfam database^33^ of protein families that currently contains 17, 929 (release 32.0) families. Domain-domain interaction data was obtained from DOMINE database ^34^ that is a collection of experimentally known and computationally predicted domain interactions. It contains 6, 634 interactions inferred from PDB entries and 21, 620 computationally predicted interactions obtained from 13 different resources.

### Confidence scoring of interologs

Since a large number of interactions are still to be discovered in biological networks, specifically in plants, so the predicted network was fragmented and contained isolated nodes. These solitary interactions were discarded and further analysis was carried out on the largest component of the TulsiPIN. Each interaction in the giant component of TulsiPIN was associated a confidence score, expressed in terms of total confidence score (*S*_*total*_). This confidence score consists of two components, one each obtained from interologous score (*S*_*ilog*_) and domain domain interaction (DDI) score which is further comprised of two terms, viz. domain interaction propensity (*S*_*dip*_) and correction for interacting-domains enrichment (*S*_*cide*_). *S*_*ilog*_ score represents the number of template plants supporting a given interaction. As PPIs in TulsiPIN are inferred from multiple template species so *S*_*ilog*_ reflects the confidence in terms of conservation of a predicted interaction. DDI score consists of two terms: first term is *S*_*dip*_, which is the ratio of reported DDI pairs in any interacting protein pair of TulsiPIN to total possible DDIs among the domains present in proteins of the same PPI pair. Second term in DDI score is *S*_*cide*_, and represents the number of reported DDI pairs between two proteins. It is introduced to account for domain interaction propensity bias resulted due to protein pairs for which a small number of Pfam domains are predicted. Domain interaction enrichment between any interacting protein pair will decrease with increase in the number of domains possessed by them while the number of reported DDI pairs between the same pair remains constant. For example, *S*_*dip*_ score for a protein pair containing one domain each (monodomain proteins) and a corresponding reported DDI in DOMINE will be equal to 1, while it is 0.25 for a pair with one reported interaction and each protein in the same pair containing two domains. Hence, an additional term in the form of (*S*_*cide*_) was introduced to account for the domain interaction enrichment bias.

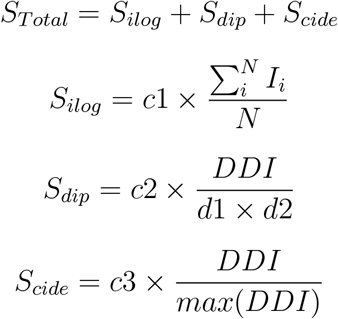

Where, *N* is the total number of species and *I*_*i*_ is equal to 1 if there exists an orthologous interaction in *O. tenuiflorum* inferred from *i*^*th*^ template plant else 0. *d*1, *d*2 are the number of domains in two proteins of an interolog, and *DDI* is the number of interacting domain pairs reported to exist in DOMINE database out of the total possible domain pairs *d*1 × *d*2 in that pair of interacting proteins. Further scaling factors of 50%, 40% and 10% in term of constants *c*1, *c*2 and *c*3 were applied to each terms of *S*_*total*_, respectively.

All the protein-protein interaction pairs having normalized score *S*_*Total*_ ≥ 0.1 were considered as high confidence interologs (HCIs) and the resulting network specific to these HCIs was termed as high confidence TulsiPIN (hc-TulsiPIN).

### Reliability assessment of TulsiPIN

Reliability of the predicted interologs was assessed using two methods i) protein colocalization similarity (PCS), ii) GO annotation based functional homogeneity index (FHI). Proteins functioning together in a particular pathway (considered to be performing similar functions) should be localized in the same cellular compartment so as to push the reaction in forward direction. This is the basic assumption of PCS method that proteins involved in an interaction should occupy the same subcompartment within the cell^35,36^. Protein localization was predicted using Deeploc^37^ keeping the default parameters. It is a deep learning based web server that predicts the location of proteins into any of the 10 cellular compartments by using benchmark experimentally annotated datasets. FHI method is based on the idea that interacting proteins perform more or less similar functions and hence GO functional annotation similarity or functional homogeneity can add an extra layer of confidence to the predicted interactions^38^. GO annotations for each protein of TulsiPIN were predicted using agriGO v2.0^39^ and were visualized using WEGO v2.0^40^ and REVIGO^41^. Jaccard coefficient was employed to estimate the functional homogeneity index for two interacting proteins that is defined as

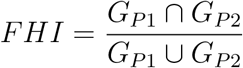

where *G*_*P1*_ ∩ *G*_*P2*_ are the GO terms shared by interacting proteins *P* 1 and *P* 2.

### Architectural properties of TulsiPIN

To identify small functional units (modules) working independently in TulsiPIN, MCODE^42^ was used. Functional homogeneity index (FHI) and colocalization similarity index (CSI) were then computed for each module. Functional homogeneity index between two interacting proteins was estimated using the method defined in previous subsection, while CSI was computed as

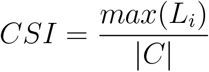

where *L*_*i*_ is the number of proteins assigned to *i*^*th*^ localization group and *|C|* is the total number of proteins for which localization could be assigned^43^. TulsiPIN proteins were also scrutinized for their involvements in various pathways using KEGG database^44^ and their potential roles as transcription factors, transcription regulators and protein kinases using iTAK package^45^.

Further, three network metrics namely degree centrality (*C*_*d*_), betweenness centrality (*C*_*b*_), and eigenvector centrality (*C*_*e*_) for each node present in the high confidence TulsiPIN were computed. Degree centrality (*C*_*d*_(*v*)) of a node *v* is the number of immediate neighbors of *v, i.e.* number of direct connections between node *v* and any other node *j* ^46^. Degree centrality for a node *v* of an undirected network can be defined as *C*_*d*_(*v*) = Σ_*j*_ *a*_*vj*_, where *a*_*vj*_ is an element of adjacency matrix **A**. Since highly connected nodes called hubs regulate a large number of other nodes, so they are more likely to be essential for maintaining and sustaining the network architecture. Also, a targeted attack on these nodes may fragment the network into several smaller components eventually making it to become nonfunctional^47^. Betweenness centrality (*C*_*b*_(*v*)) of a node *v* is computed as

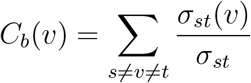

where *σ*_*st*_ is the total number of paths existing between any two nodes *s*and*t* while *σ*_*st*_(*v*) is the number of paths passing through *v* ^46^. High value of *C*_*b*_(*v*) indicates relative importance of nodes connecting different communities than those which are central to them^48^. Eigenvector centrality (*C*_*e*_) prioritizes nodes in a network by considering the number of connections possessed by their immediate neighbors along with their own degrees. It is based on a simple idea that any node’s importance in the network depends upon both its own degree as well as the degrees of its neighbors^49^. Additionally, two global network measures, namely, average path length (*ℓ*) and clustering coefficient were calculated for the TulsiPIN and for its random realizations obtained as per the methods described in the next subsection. Average path length or average geodesic distance is computed as^50^

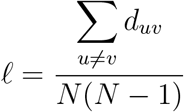

where *d*_*uv*_ is the distance between nodes *u* and *v* while *N* represents network order. Clustering coefficient (*C*(*v*)) of a node is a measure of its connectedness and can be computed as

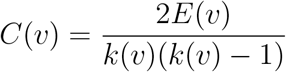

where *k*(*v*) is the degree of node *v* and *E*(*v*) is the number of edges existing between neighbors of *v* ^50^.

### Identification of statistically significant bottleneck proteins

For identifying the key nodes in hc-TulsiPIN, we first constructed two ensembles each consisting of 10, 000 corresponding random realizations. In the first ensemble, random networks were generated using *G*_*n,m*_ type Erdős-Rényi (ER) model^51^ that preserve the network size and average degree of the hc-TulsiPIN. While in the second ensemble, scale free (SF) random graphs were generated using an extended version of Barabási-Albert (BA) model^52,53^. These SF type random networks consist of identical number of nodes while edges differing within the range of 1% from that of hc-TulsiPIN. For that, initially a set of *i* isolated nodes was selected such that *i* ≤ *k* ≤ *i* + 1 where 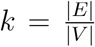. Then each new node was introduced to the network with *i* or *i* + 1 new edges having probability *i* + 1 − *k* and *k* − *i* respectively. Addition of new nodes follows preferential attachment, having more likelihood of connecting with high degree nodes than to low degree ones. Statistically significant central nodes were identified by comparing the hc-TulsiPIN with the ER and BA type random networks of previously designed ensembles in terms of three network metrics, viz. *C*_*d*_, *C*_*b*_ and *C*_*e*_. Statistical significance for any node *v* of hc-TulsiPIN was estimated using *z-score* that is defined as

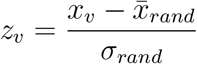

where, *x*_*v*_ is the value of a network metric of the *v*^*th*^ node in the hc-TulsiPIN, and 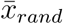, *σ*_*rand*_ respectively are the mean and standard deviation of the selected ensemble. All the nodes having *p <* 0.01 were considered statistically significant. Finally, the nodes that were predicted statistically significant in comparison to both the random models constitute the set of vital proteins in TulsiPIN.

## Results and discussion

### *O. tenuiflourum*’s protein-protein interaction network (TulsiPIN)

In any living cell, most of the biological processes are governed by small molecules which usually function in concert to accomplice and give rise to multitude of complex cellular behaviors. Proteins, for example, interact transiently with other proteins forming macromolecular complexes to perform specific functions. So decoding a protein’s function demands identification of the functional partners (permanent or transient) it physically interacts with. Corollary of this has been implemented in assigning functions to uncharacterized proteins from their interacting partners with known functions^54^. Despite huge advancement in high-throughput technologies, PPI information of plants is sparingly and sparsely available. Owing to PPI data scarcity and high cost, time, labor *etc.* incurred on experimental determination of PPIs, computational approaches which employ already existing information from other datasets to generate new information for the concerned organism have attracted researchers from past few decades. One such predictive approach is called interolog method, that assumes structural (architectural) and functional conservation of protein interactions across species during the course of evolution. The method exploits this cross-species conservation to transfer already known interactions from template species to the desired organism. This inferred interactome is thus an outcome of orthology between respective proteins in query and template proteomes, resulting in prediction of interacting protein pairs called interologs ^55^. In the present study *O. tenuiflorum*’s proteins possessing “one-to-one” orthologous relationships with 49 template plants were identified using homology search tool BLASTp. Two proteins were considered to be mutually interacting if there exists at least one interaction between their corresponding orthologous proteins in any of the template interactomes (see Methods Section). Out of the 49 template plants, 36 plants were found to have at least one orthologous interaction to *O. tenuiflorum* (see Table 1). This resulted in an interologous protein interaction network (TulsiPIN) consisting of 13, 660 nodes/proteins and 327, 409 unique binary pairs of interactions. The largest proportion of interologous interactions is obtained from *Erythranthe guttata* followed by *Gossypium raimondii*, *Amborella trichopoda*, *Arabidopsis thaliana* etc. Table 1 lists the template species along with interactions inferred from each template. A substantial decrease in the proportion of unique interactions contributed by the successive template plants reflects the exhaustiveness of potential PPIs in *O. tenuiflorum* evaluated for the construction of TulsiPIN.

**Table 1:**
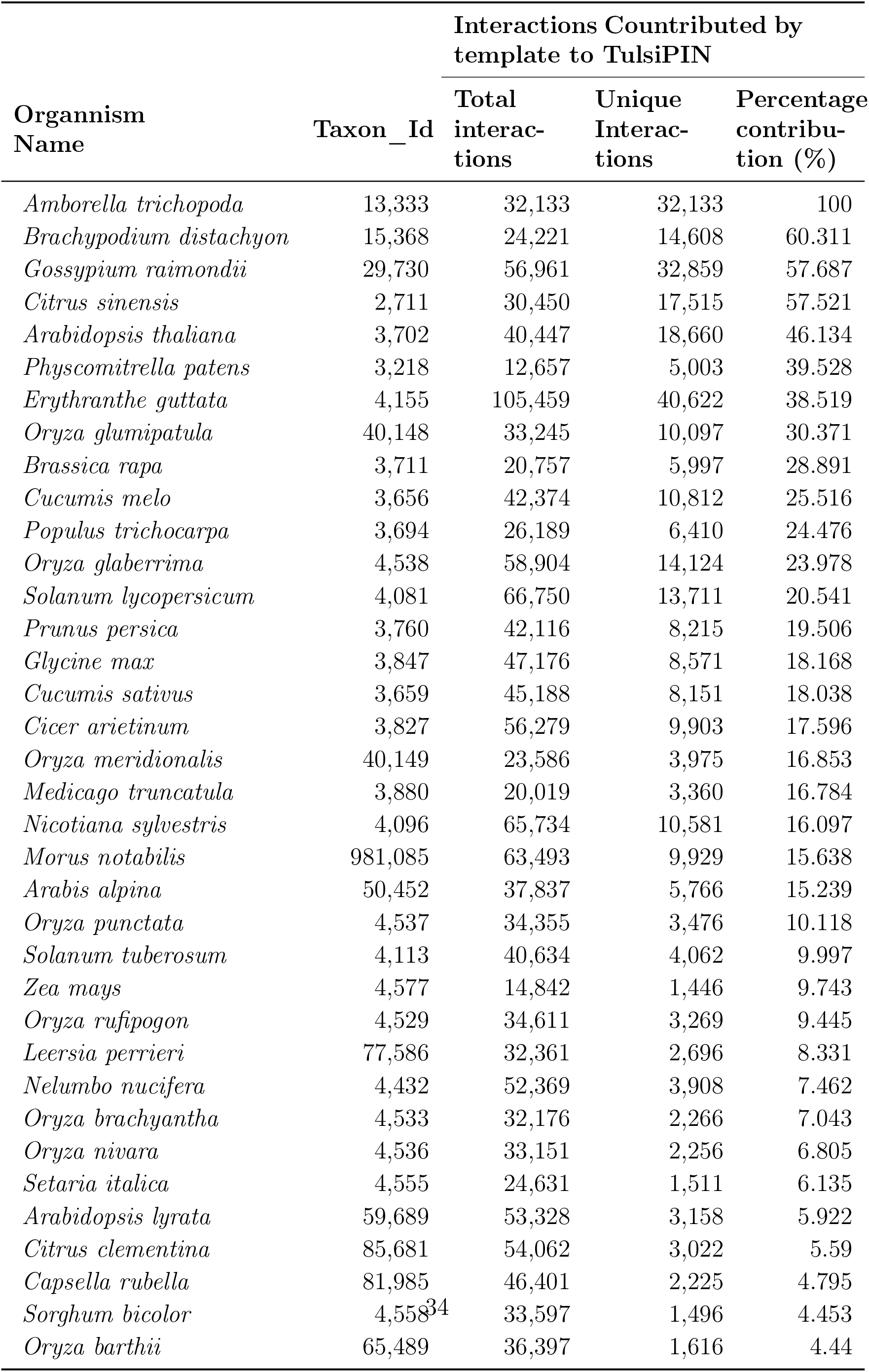
Proportion of unique and total interactions contributed by individual template plant in predicted TulsiPIN

## Construction of a high confidence TulsiPIN (hc-TulsiPIN)

Since the TulsiPIN is a predicted network consisting of interactions that are inferred from different plant templates, we evaluate the confidence of each interaction by associating a confidence score (*S*_*total*_). In the following, we detail various steps involved in obtaining the *S*_*total*_ for each interaction.

### Domain-domain interaction based assessment

First of all, we enumerated the potential domain-domain interactions underlying PPIs of TulsiPIN from the collective information of interacting domains present in DOMINE database. Structural studies on PPIs in past few decades have revealed that different proteins interact through small structural subunits called domains. Also, a large number of proteins (approximately two third in eukaryotes) are known to be multidomain *i.e.* consisting of more than one domains^30^. Since different domains possess unique structures and biological functions, so they are believed to have been designed and evolved to interact with specific domains^56^. An average PPI hence is not determined by all the domains possessed by proteins participating in the interaction rather by a selected subset only^32,57^. Thus, putative domain-domain interactions (DDIs) supporting the predicted PPIs were identified by leveraging exhaustively collected DDI information stored in DOMINE database. First, domain architectures of all the proteins included in TulsiPIN were obtained by querying them against Pfam database. The search resulted in 12, 082 (*≈* 94% of total 13, 660) proteins containing one or more domains. Then, the non-redundant interactions between every possible domain-pair among the domains contained in an interolog of TulsiPIN were searched in DOMINE. Out of 12, 082 proteins having been assigned at least one domain, 5, 757 proteins (*≈* 42% of total TulsiPIN and *≈* 48% of the proteins predicted to have at least one Pfam domain) were indicated by DOMINE to participate in domain-domain interactions. Accordingly, domain interactions supported 48, 645 (*≈* 15%) of the total predicted PPIs in TulsiPIN.

### Ranking of interologs

Further, each PPI of TulsiPIN was assigned a confidence score (*S*_*total*_) which ascertains the reliability of predicted interactions. This confidence score consists of two components,one each obtained from interolog and DDI methods. Scores obtained from each method were assigned equal weights (50%). Since proteins interact with each other through specific domains to carry out specialized processes vital for growth and development of an organism, interacting domains information can play crucial roles to apprehend the complex mechanisms underneath protein interactions as well as to serve as basis to mark the strength of predicted interactions. We ranked all the interologs using a confidence score *S*_*total*_, where magnitude of score reveals the strength of interaction. Interactions falling in the lowest 5% of *S*_*total*_ score, corresponding to *S*_*ilog*_ *<* 3 and *S*_*dip*_ = 0, were filtered out as low confidence interologs (LCI) as they are marginally supported by *S*_*ilog*_ obtained from few template plants and have no support from domain domain interaction score (supplementary Table S2). We obtained 184, 852 interactions which fall under the LCI category and removed from the network. Then the *S*_*total*_ score was normalized between [0, 1] using min-max scaling and interologs falling in the lowest 10% of normalized *S*_*total*_ (i.e. having normalized score between 0 and 0.1) were considered as moderate confidence interologs (MCI). A total of 46, 917 protein pairs were found to have normalized *S*_*total*_ score less than 0.1 and were tagged with moderate confidence interologs (supplementary Table S2). On the other hand, interactions scoring more than or equal to 0.1 of normalized *S*_*total*_ were considered as high confidence interologs (HCI). A total of 95, 605 interacting pair among 7, 845 proteins were found to have normalized *S*_*total*_ *≥* 0.1 (see Table 2). However, there were some isolated interactions present in this high confidence network so we considered only the largest component that is comprised of 7, 719 nodes and 95, 532 interactions (Figure 4a) and is termed as high confidence TulsiPIN (hc-TulsiPIN).

**Table 2:** Interaction details of TulsiPIN

| Interaction Type | Complete Network |  | Confidence Score | Largest Component |  |
| --- | --- | --- | --- | --- | --- |
|  | Interactions | Proteins |  | Interactions | Proteins |
| TulsiPIN | 327,409 | 13,660 | - | 327,374 | 13,604 |
| HCI | 95,605 | 7,845 | 0.1 - 1.0 | 95,532 | 7,719 |
| MCI | 46,917 | 7,390 | 0.0 - 0.1 | - | - |
| LCI | 184,852 | 13,121 | - | - | - |

This scoring schema represents an intuitive classification that instead of outrightly categorizing interologs into different classes, combines the interologous and domain-domain interaction approaches to classify them into three categories. The proposed methodology could successfully discriminate between pairs which were supported by a large number of template plants but not having domain-domain interactions and assigned them lower weights. Interologs supported by as large as 7 template species were not considered in high confidence network because these interactions were either having no or very small contribution of *S*_*dip*_ and *S*_*cide*_ ((supplementary Table S2)). Similarly, large number of pairs which were supported by as many as 30 template plants are ranked low because they are supported by interolog method only. Contrary, there were a large number of protein pairs which were ranked very high in the network due to having substantial support from the domain-domain interactions, although they were supported by fewer than 10 template plants. Similarly, this methodology has very carefully ranked multidomain pairs higher over monodomain pairs when both were supported by same number of template plants.

### Reliability assessment of hc-TulsiPIN’s interologs

hc-TulsiPIN, as reported in this study, is a homology based protein-protein inetraction network constructed from 36 different plant templates. Therefore, we attempt to assess the confidence of reported PPIs in this network using available functional information.

#### Functional homogeneity based assessment

Functional coverage of predicted hc-TulsiPIN was then assessed by gene ontology (GO) analysis. By means of GO annotation, each protein of hc-TulsiPIN was subjected to assignment of GO terms. A total of 9, 091 GO terms were successfully associated with 5, 278 proteins through GO annotation. Among these, 3, 294 GO terms were classified into 17 categories of “Biological Processes” in which ‘metabolic process and cellular process’ categories were found to be most enriched followed by ‘localization’ and ‘biological regulation’. 4, 632 GO terms were associated with 8 categories of “Molecular Functions” with abundance of “binding and catalytic activity”, “structural molecule activity” followed by “transporter activity”, and “molecular function regulator” that reveals coherence between the biological processes in which hc-TulsiPIN’s proteins are involved and the molecular functions they perform. Remaining 1, 165 GO terms were categorized into 9 classes of “Cellular Components” with “cell part” and “cell” being highly enriched followed by “organelle” and “protein-containing complex” (Fig 2). Assignment of gene ontology (GO) terms to several proteins reveals that the node proteins of hc-TulsiPIN are belonging to diverse gene families.

**Figure 2:**
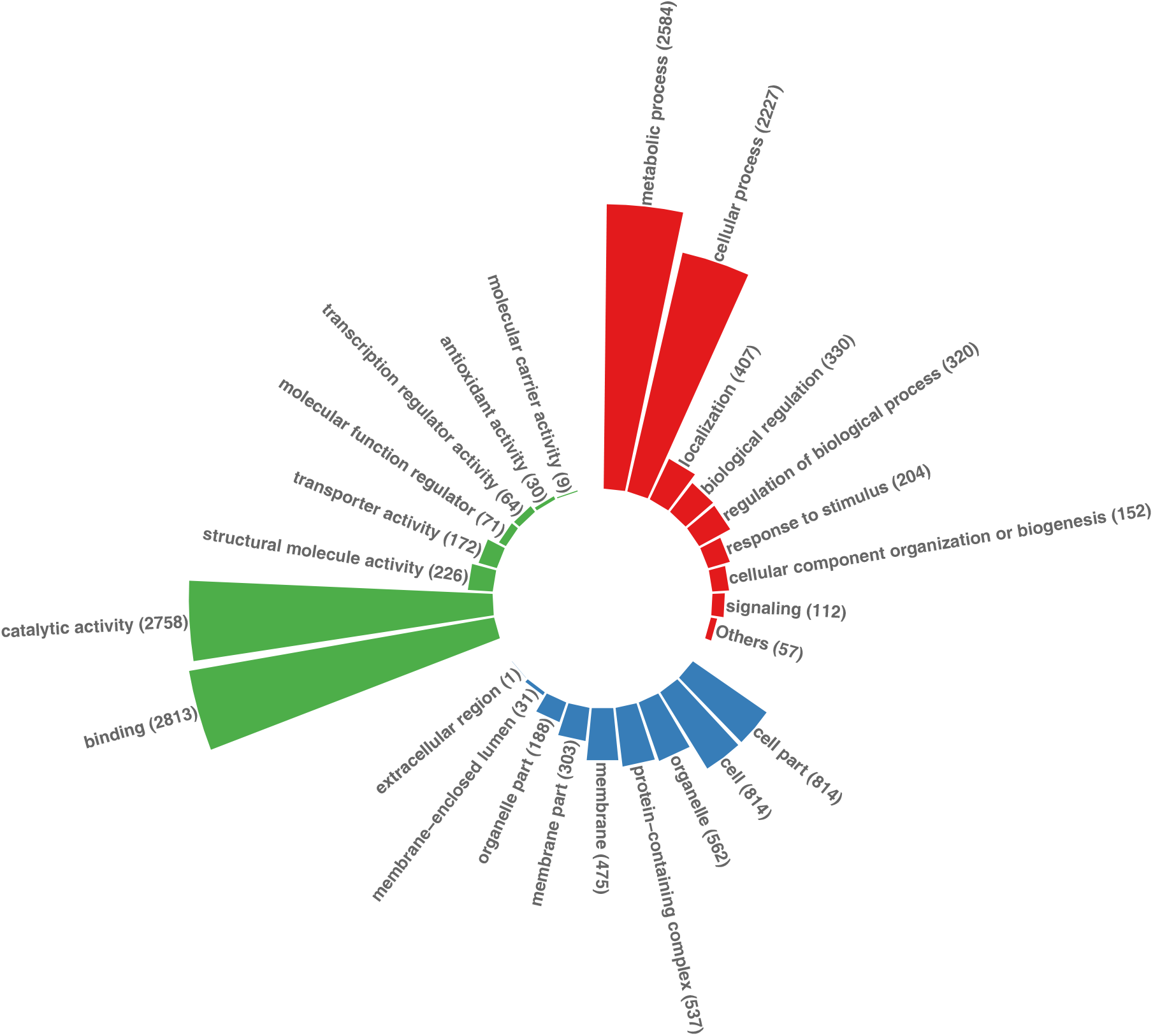
GO based functional enrichment of 7, 719 proteins involved in hc-TulsiPIN grouped according to three functional ontologies namely, cellular component (3, 725 proteins colored in blue), molecular function (6, 143 proteins colored in green) and biological process (6, 393 proteins colored in red).

Reliability of predicted network was further assessed by computing the functional homogeneity of each PPI pair and comparing them with randomly placed proteins together in an interaction. A total of 52, 032 protein pairs were obtained for which GO terms could be assigned in the high confidence hc-TulsiPIN. Average FHI obtained for hc-TulsiPIN was approximately 0.218. Although, maximum and minimum number of PPIs for which GO terms were assigned in 10, 000 G(n, m) type ER random networks were 45, 220 and 44, 094, however, their corresponding FHI values were 0.029 and 0.031 respectively (supplementary Table S1). Average FHI obtained for hc-TulsiPIN was significantly high (7 times) than average FHI (0.031) obtained from random ensemble. This high value of FHI for PPIs in high confidence hc-TulsiPIN reveals strong coherence in function between connected protein pairs than those placed randomly together in an interaction. Moreover, despite comparable number of PPIs for which GO terms were assigned in both real and corresponding random ensemble, PPIs predicted in hc-TulsiPIN are found to be performing similar functions compared to PPI pairs of the random networks.

#### Protein colocalization based assessment

Subcellular localization of all the 7, 719 hc-TulsiPIN proteins were successfully predicted using Deeploc (v1.0) into ten subcellular locations. Maximum number of proteins were predicted to be cytoplasmic (3, 001) followed by nuclear and mitochondrial having 1205 and 1016 proteins respectively. Further the subcompartmental information was mapped to PPIs of high confidence hc-TulsiPIN. A total of 22, 527 PPI pairs were predicted as cytoplasmic while 6, 913 as nuclear. 2, 952 PPIs were predicted between proteins present in plastids while 2, 211 PPIs were between mitochondiral proteins. To add another layer of confidence to the high confidence network, we examined the subcellular locations of proteins participating in PPIs in comparison to those of the ER type random networks. Figure 3 depicts the distribution of frequencies of PPIs in different subcellular compartments in hc-TulsiPIN (dots with empty centres) alongside randomly linked PPIs. Frequencies of random PPIs are averages over ensemble of 10, 000 G(n,m) type ER networks and bars represent the range of frequencies. Results revealed a complete deviation of PPI frequencies in hc-TulsiPIN from the random ones, with frequencies of hc-TulsiPIN being significantly high for most of the locations. Functional analysis of the high confidence hc-TulsiPIN *i.e.* reveals that most of the proteins involved in an interaction are residing in the same subcellular compartments and are performing similar functions than expected by random chance.

**Figure 3:**
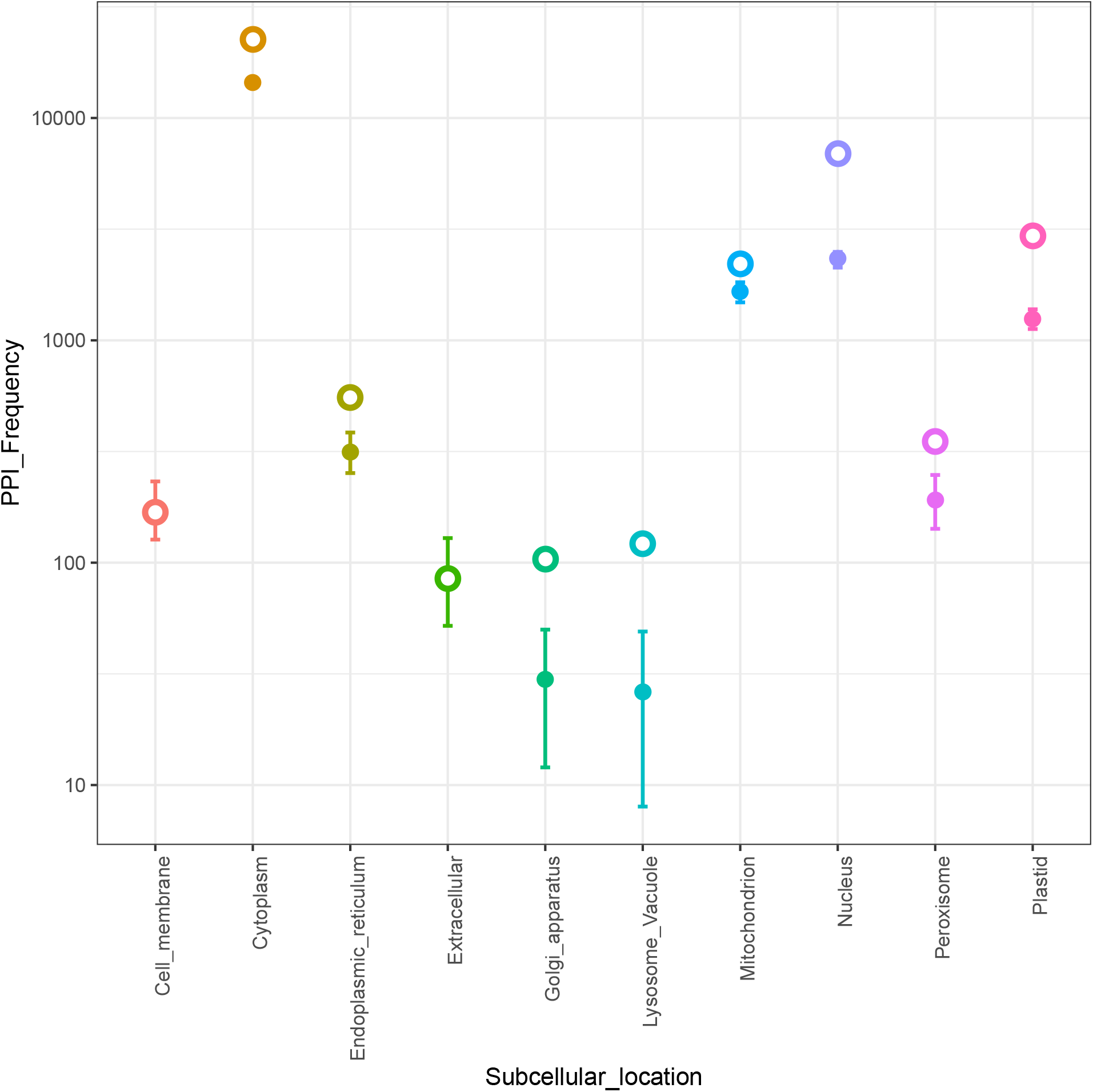
Comparison of PPI frequencies of hc-TulsiPIN (represented by open circles) with the corresponding PPI frequencies in 10, 000 realizations of ER type random networks (represented by filled circles with range bars) in 10 sub-cellular compartments.

### Topological characteristics of hc-TulsiPIN

In this section, we report the computational results of some of the well established topological properties attributed to biological networks for hc-TulsiPIN. Three global network metrics including degree centrality, betweenness centrality and eigenvector centrality were computed for hc-TulsiPIN. We also investigated two global network properties for hc-TulsiPIN, which are small worldness and scale-free degree distribution (see Figure 4b). The degree distribution of hc-TulsiPIN when plotted on a double logarithmic scale follows a power law (*P* (*k*) ~ *k*^*−γ*^) characteristic to scale free networks^58^, with exponent *γ* having value *−*1.826 (see Table 3). Another very important global property of biological networks is the small world property that has an important role in spreading the information throughout the network. Information flows quickly in very short time to a large number of nodes in networks possessing small world property^59^. Networks with low average path length (*ℓ*) and high clustering coefficient (*C*) compared to random networks of comparative dimensions *i.e*. *ℓ*_*smallworld*_ ≥ *ℓ*_*random*_ and *C*_*smallworld*_ » *C*_*random*_ are considered to have small world architecture^59^. hc-TulsiPIN was subjected for the estimation of both these global metrics and the values obtained were found to have an explicit deviation from those obtained for the random networks (Table 3). Average path length (*ℓ*) of hc-TulsiPIN was found to be 4.014 which is slightly higher than the one obtained from random networks (3.06) while the average clustering coefficient (*C*) obtained for hc-TulsiPIN (0.310) was much higher (more than 100 fold) than the corresponding random counterpart (0.003) obtained by averaging over 10, 000 similar ER networks (Table 3).

**Table 3:** Global network properties of hc-TulsiPIN along with corresponding averages of 10,000 ER and SF random networks

| Network Dimensions and Properties | Network Types |  |  |
| --- | --- | --- | --- |
|  | ER | SF (PCJ) | hc-TulsiPIN |
| Order ($n$) | 7,719 | 7,719 | 7,719 |
| Size ($m$) | 95,532 | 95,383 | 95,532 |
| Power law exponent ($\gamma$) | 7.881 | 1.621 | 1.826 |
| Clustering coefficient ($C$) | 0.003 | 0.0157 | 0.310 |
| Average path length ($\ell$) | 3.068 | 2.887 | 4.014 |

**Figure 4:**
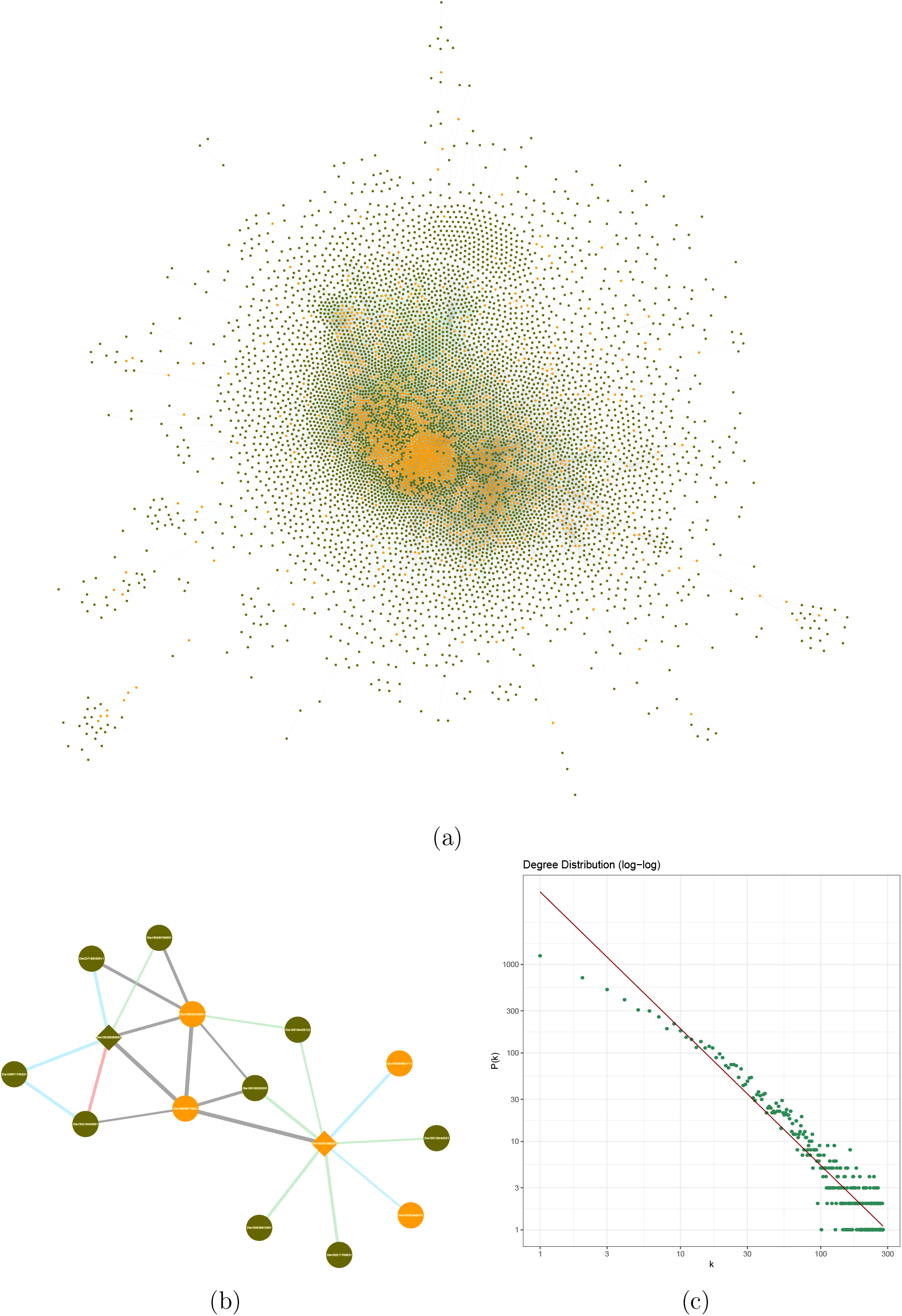
Topology of hc-TulsiPIN: (a) Complete hc-TulsiPIN having 7, 719 nodes and 95, 532 interactions. Key nodes are highlighted in orange color. Edge widths are scaled to the associated confidence score. Each edge is colored according to the number of reliability measures supporting that interaction: pink - 3, cyan - 2, green - 1 and grey - no support. (b) A subnetwork of two proteins, namely, Ote100256650051 and Ote100091090201 (both being the geraniol dehydrogenases, and represented as diamonds) with their first degree interactors highlighting the coloring schema and relative strength of various edges. (c) Degree distribution of hc-TulsiPIN with power law fit having exponent *−*1.826.

### Identification of key proteins

Although there are very few influential nodes in a network, however, their influence in terms of function or catastrophic failure spreads across entire network in a quick time. Many centrality measures in the past few decades have been proposed to identify these topologically and hence functionally important bottlenecks in a biological network^60^. However, identifying these key proteins is a hard job due to few reasons i) Different centrality measures are designed to address different architectural aspects of a network and therefore capture the picture of a network in terms of selected features only and emphasize the vitality of nodes only in a particular aspect. For example, degree centrality is a connectivity based metric so it enumerates the importance of a node in terms of connectivity only, on the other hand betweenness centrality assigns higher rank to nodes connecting two large communities^60^. ii) A centrality metric ranking a node higher does not always guarantees its essentiality and its corollary that nodes ranked lower are non-vital is not always true. Due to reasons like unavailability of complete interactomes, reliability issues will always spur on quantifying the importance of nodes. It is almost impossible to design an unified measure that can quantify a node’s importance in every possible situation.

Towards addressing these issues, we propose a hybrid approach for identifying key proteins in the TulsiPIN. In our method, a protein is said to be a key protein of TulsiPIN if it satisfies following two criteria: (i) It must appear in the hc-TulsiPIN, as these proteins are considered to have a more reliable interaction than the other proteins of complete TulsiPIN. (ii) It must have a high value (in terms of *z-score*) of any of the three topological metrics, viz. degree, betweenness and eigenvector centralities in the hc-TulsiPIN, in comparison with both the random network ensembles. For that, we first associate statistical significance with all the proteins of hc-TulsiPIN. Six *z-scores* were computed for each node by comparing its three network metrics values with both the random ensemble of 10, 000 networks of ER and BA type models. For details on the construction of these ensembles, please visit the Methods Section. Against each of the random ensemble, a set of statistically significant proteins was designed consisting of proteins having *p-values ≤* 0.01 for any of the three network metrics. Finally, proteins that were found to be statistically significant in both the sets were considered as the key proteins. This resulted in identification of 1, 625 key candidates in the hc-TulsiPIN. Supplementary Table S3 enlists all the important nodes along with putative functions they perform in *O. tenuiflorum*.

### Modular architecture of high confidence TulsiPIN

After enumerating various descriptive statistics of TusliPIN, an attempt was made to establish a relationship between network organization and its function by identifying specific subset of proteins working in concert to bring about common function called modules. Extraction of functional modules like protein complexes^61^ or pathway proteins^62^ from global PPI networks is an important step towards knowledge discovery. A total of 157 high density subnetworks including 2, 459 nodes were extracted from TulsiPIN by leveraging molecular complex detection (MCODE) algorithm. Owing to the facts that (i) two proteins can interact only when they co-localize in same cellular compartment and (ii) proteins participating in an interaction are more likely to be performing same function, we estimated the functional content of hc-TulsiPIN in terms of functional homogeneity index (FHI) and colocalization similarity index (CSI) for all the identified modules (supplementary Table S2). Functional homogeneity index, in terms of JC, was computed for each module (supplementary Table S4). A total of 71 modules are found to have FHI values higher than than the average FHI of hc-TulsiPIN. Further, a total of 15 modules having more than ten proteins that were assigned at least one GO term and FHI greater than average FHI (0.218) of hc-TulsiPIN are selected for colocalization similarity assessment and pathway enrichment analysis. Among the selected 15 modules, the highest functional homogeneity index is obtained for module_22 (0.87) followed by module_1, and module_77 with FHI scores 0.53 and 0.47, respectively. High FHI indicates that the interacting proteins in a module are involved in or performing similar functions. Further, 65% modules (102) from a total of 157 are found to have colocalization similarity index (CSI) *≥* 0.5. CSI for the selected 15 modules is ranging between 0.818 *−* 0.266 which indicates most of the proteins in a module colocalize with in the same subcellular location. This fact has been further confirmed when we mapped the pathways in which proteins of each of these 15 modules are functioning. Most of the modules reveal same pathways as those predicted by gene ontology terms. Supplementary Table S2 lists the selected 15 modules along with most enriched GO terms and subcellular localization where most proteins are localized in. Moreover, most of the modules identified are related to genetic information processing, transport, lipid and crbohydrate metabolism.

### Pathway analysis of network proteins

To find the important regulatory pathways in which TulsiPIN proteins are functioning, we used KEGG database, and found that a total of 5, 665 proteins with 2, 005 unique KO ids encoding 298 pathways which are further classified into five major categories, viz. Metabolism, Genetic information processing, Environmental information processing, Cellular processes and Organismal systems as shown in Fig 5. Further, metabolism category is divided into 11 subcategories in which maximum number of proteins are reported from “carbohydrate metabolism” (1432), subsequently followed by “amino acid metabolism” (879) and “lipid metabolism” (654). Under genetic information processing, translation (758) is most enriched followed by folding, sorting and degradation containing 676 proteins. All the proteins involved in these pathways are very important as they are required by the plant for their basic processes like growth, development and response to different pathological conditions etc^63–65^. Two classes including “signal transduction” (1, 983) and “membrane transport” (39) are found highly enriched under environmental information processing subcategory. These are the proteins that work under fluctuating environmental conditions and are crucial for the plant to adapt such adverse conditions^66^. In “cellular processes” subcategory “cell growth and death”, “transport and catabolism” were among the most enriched classes containing 817 and 799 proteins respectively. Finally, in the “organismal systems” subcategory 10 systems are predicted among which “endocrine system” (661) was found to be most enriched followed by “immune system” (520), “nervous system” (415) etc. We further mapped the key proteins to pathways and found that from a total of 5, 665 proteins predicted to function in different pathways, 4, 184 belong to hc-TulsiPIN. Most of the proteins are related to “Translation” (283), followed by “Folding sorting and degradation” (181), “Transport and catabolism” (151). Among these 4, 184 proteins, 1, 155 are key proteins and most of the key proteins predicted by our methodology are predicted to be functioning in pathways related to “genetic information processing” like “translation”, “transcription”, “folding sorting and degradation”, “replication” *etc.*.

**Figure 5:**
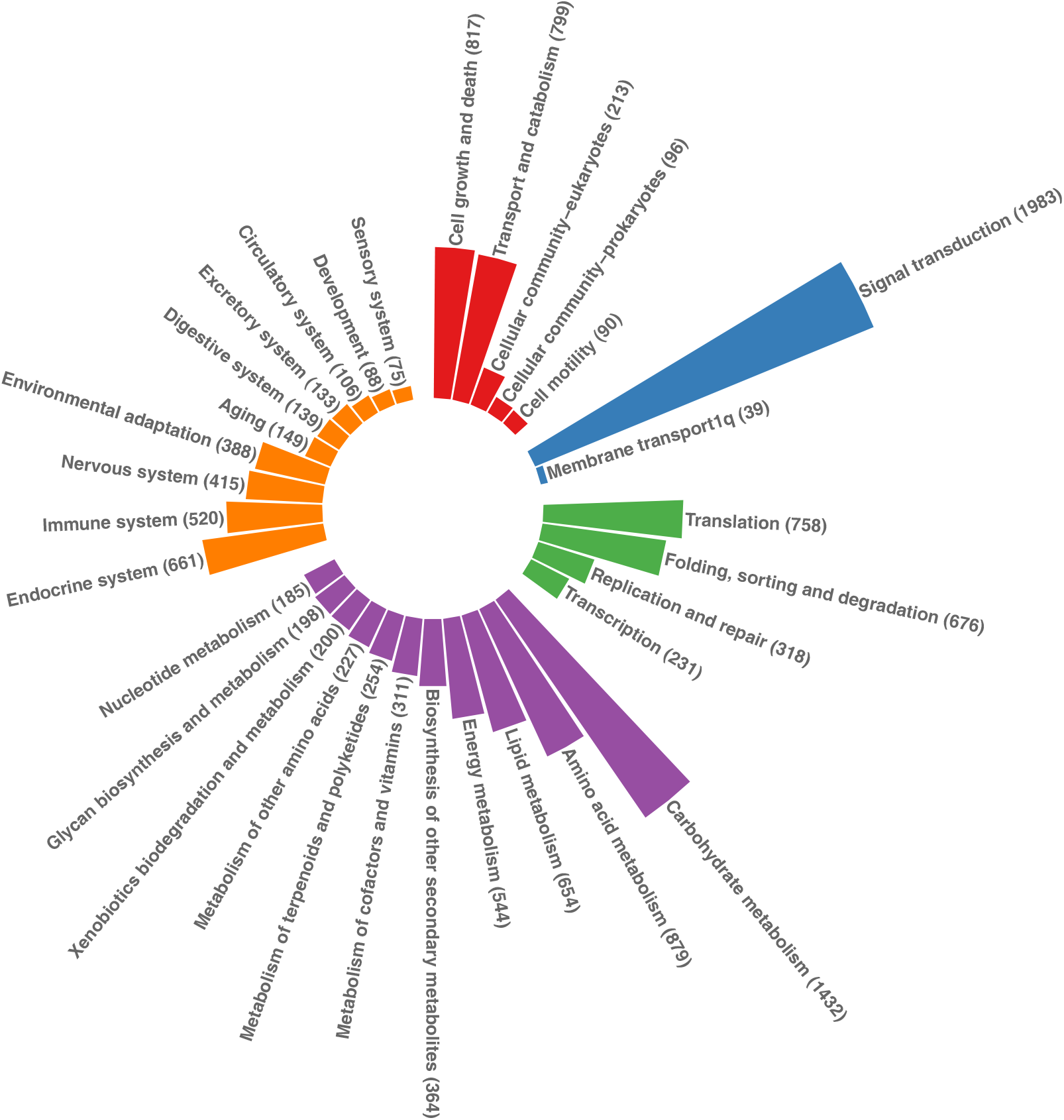
Pathways enrichment of 7, 719 proteins involved in hc-TulsiPIN grouped by major biological processes, viz. metabolism (5,248 proteins colored in purple), genetic information and processing (1,983 proteins colored in green), environmental information processing (2,022 proteins colored in blue), cellular processes (2,015 proteins colored in red) and organismal system (2,674 proteins colored in orange).

### Identification of transcription factors (TFs), transcription regulators (TRs) and protein kinases (PKs)

The whole proteome of *O. tenuiflorum* was subjected to iTAK package as input to mine three types of regulatory proteins viz. TFs, TRs and PKs. From all the proteins of complete TulsiPIN 467 are identified as TFs, 243 as TRs and 666 as protein kinases (PKs). Further, these 243 TRs are categorized into 24 different families while 467 TFs are classified into 60 families. Out of 467 TFs, 134 proteins are encompassed by top five families namely MYBrelated, bHLH, NAC, bZIP and GRAS listed in descending order of number of proteins present in each family Fig 6. MYB and bHLH families are most populated TF families containing 31 proteins each while bZIP (25), NAC (24), GRAS (23) are the subsequent categories with number of *O. tenuiflorum* proteins predicted as belonging to the respective family given in parenthesis (Supplementary Table S5). The subnetwork of TFs in complete TulsiPIN consists of 3, 083 nodes (467 TFs and 2, 616 first neighbors) and 5, 733 unique interaction pairs. TF Ote100097240021 that is predicted to be a member of MYB family, is most connected protein among all the transcription factors. It shares 724 direct links with other proteins, and has high betweenness centrality (*≈* 0.5) that reflects its crucial role as a mediator of information spread between its neighbors. MYB family of TFs plays crucial roles in large number of plant processes namely biotic (pathogenic) and abiotic stress response, primary and secondary metabolism^67^, disease resistance, cell fate determination, plant development which includes anther development, root; shoot; meristem development etc.^68^. All the members of family contain a helix-turn-helix (HTH) fold that render them the ability to bind DNA molecules. Subsequent high degree TFs include Ote100006770191 (192), Ote100032970041 (171), Ote100045400121 (162), Ote100099560093 (160) belonging to C2H2, S1Fa-like, NF-YB and C2H2 TF families respectively. Out of 467 TFs, 147 could successfully be mapped in the hc-TulsiPIN, among them 17 are the key proteins.

**Figure 6:**
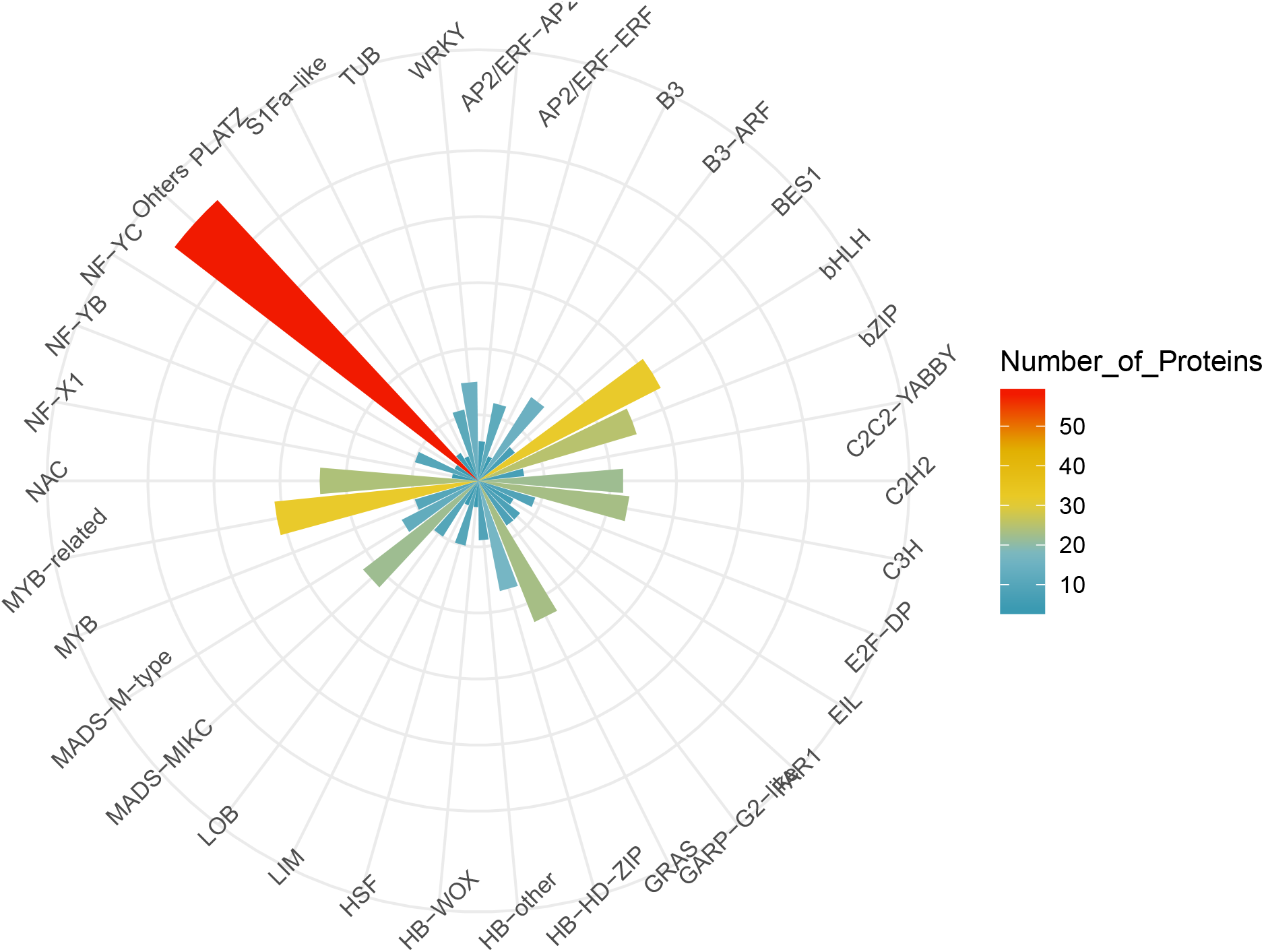
Distribution of hc-TulsiPIN’s proteins in 60 different categories of transcription factors. Length and color intensity of each sector represent the number of proteins in individual category.

Transcription regulators (TRs) are different from TFs in that they indirectly regulate transcription by limiting or exposing the chromatin to TFs (cis-elemetns). From a total of 710 regulatory proteins 243 are categorized in 24 different families of TRs. Top five families of TRs are found to be SNF2, SET, AUX/IAA, TRAF and GNAT containing 39, 25, 21, 18 and 17 proteins respectively (Supplementary Table S5). The transcription regulators subnetwork of *O. tenuiflorum* consists of 243 TRs and 1, 741 other proteins which interact among themselves and other proteins in the complete TulsiPIN to give rise to 5, 952 binary interactions (Figure 7). Ote100098110091 and Ote100109180281, predicted members of HMG and GNAT families, are found to have highest number of interactions (208 and 207) with other proteins followed by Ote100247460021 (166), Ote100011800041 (127), Ote100031230081 (125). Both Ote100247460021 and Ote100031230081 are predicted as members of SNF2 family while Ote100011800041 is predicted as SET family TR. Members of SNF2 family are involved in important process like chromatin remodelling that limits DNA accessibility, regulation of DNA methylation that has a very important role in flowering plants^69^. Among 243 TRs a total of 179 proteins are found to be belonging to hc-TulsiPIN while 21 of them are the key proteins.

**Figure 7:**
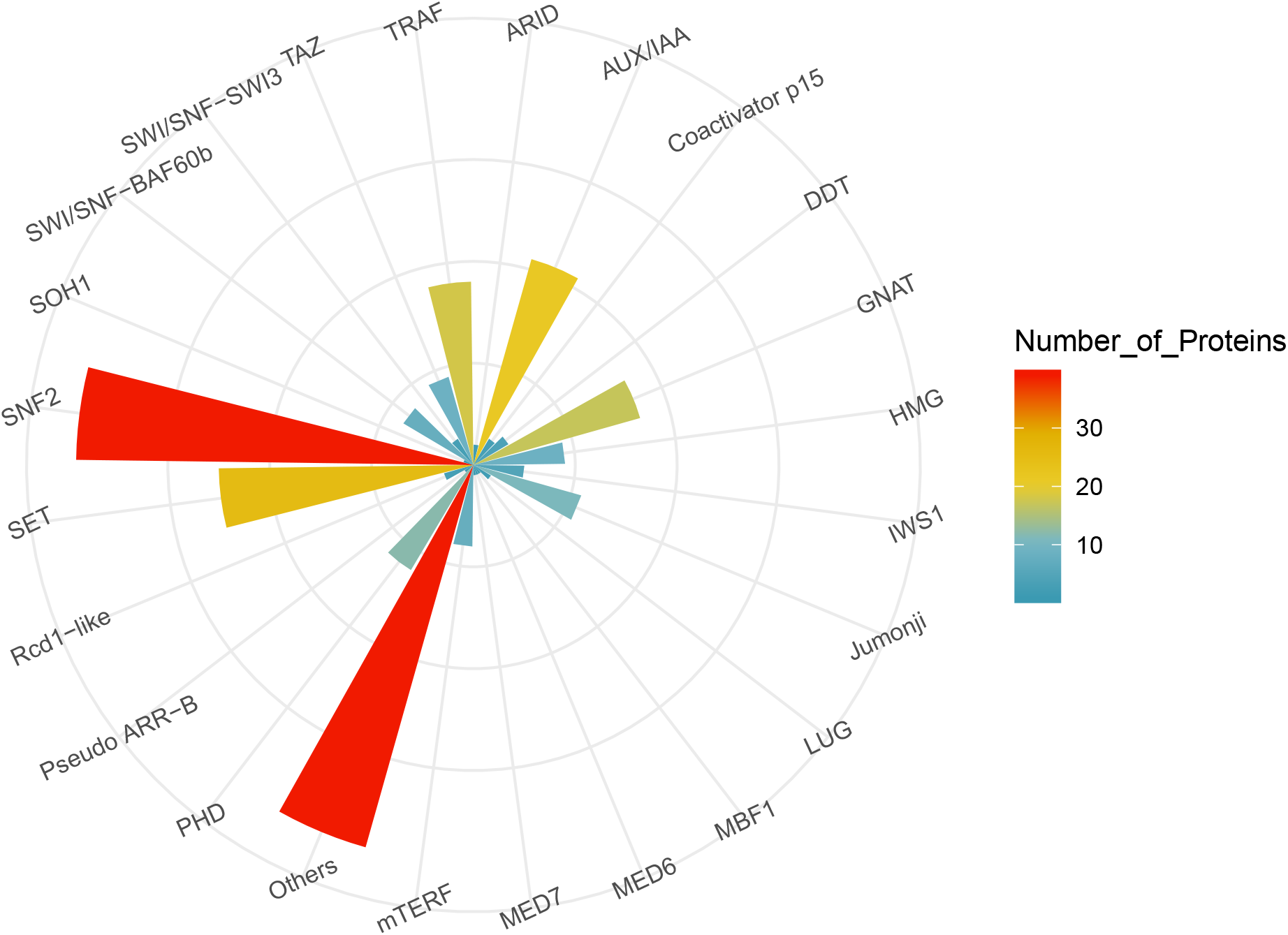
Distribution of hc-TulsiPIN’s proteins in 24 different categories of transcription regulators. Length and color intensity of each sector represent the number of proteins in individual category.

We obtained 666 protein kinases classified into nine major categories viz. RLK-Pelle, CMGC, CAMK, AGC, STE, CK1, Plant_Specific, TLK and others, from a total of 13, 604 *O. tenuiflorum* proteins. 319 proteins are predicted as RLK-Pelle type of kinases, 86 were identified as CMGC type and 85 as CAMK type of PKs (Supplementary Table S5). From a total of 666 protein kinases, 528 proteins are successfully mapped to hc-TulsiPIN in which 57 are found to be key proteins. In the complete TulsiPIN, these 666 PKs are found interacting with 2, 829 other proteins and constitute among them 9, 468 interactions (Figure 8). Top five proteins on the basis of degree in the protein kinase subnetwork are Ote100055550081 (1, 435), Ote100073170111 (289), Ote100158690021 (261), Ote100025130071 (248), Ote100132560011 (203). Further mapping of key proteins extracted from hc-TulsiPIN to the predicted kinases revealed that 27 of the bottlenecks are kinases.

**Figure 8:**
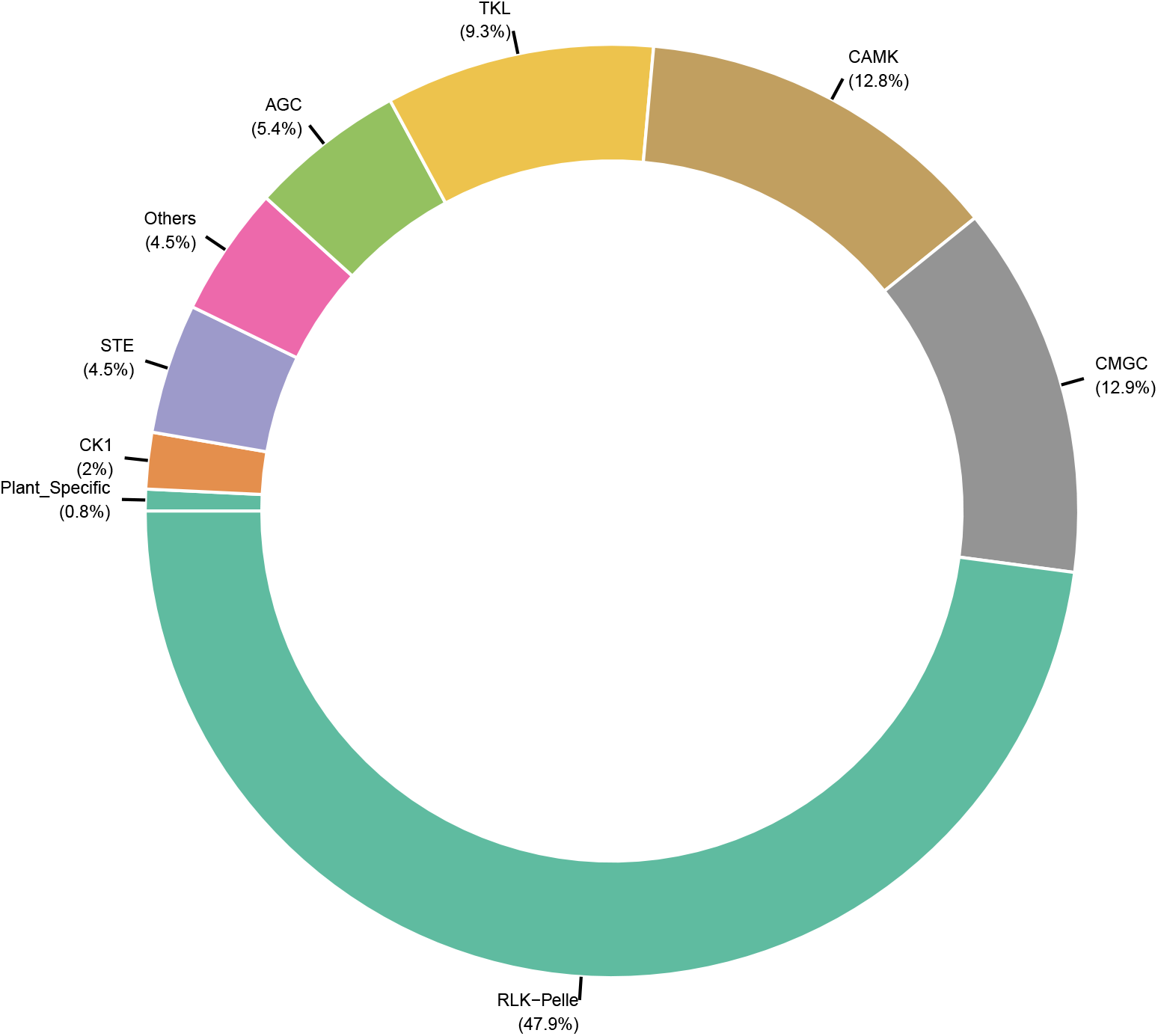
Distribution of hc-TulsiPIN’s proteins in 9 different categories of protein kinases.

### Regulatory network of proteins involved in selected metabolite biosynthetic pathways

Metabolism encompasses all those processes vital for a living being while all the metabolic reactions either catabolic or anabolic are catalyzed by enzymes (biocatalysts). As the medicinal values of *O. tenuiflorum* are attributed to the metabolites it produce, so delineating the complex pattern of interplay between different proteins may provide important insights into biosynthetic pathways of these metabolites. Here, to represent a holistic view of how genes or proteins involved in important secondary metabolites biosynthetic pathways, 181 genes encoding 14 pathways were obtained form TulsiDB. These 181 guide proteins could be successfully mapped to 76 proteins in TulsiPIN encompassing 10 pathways. Collectively 670 nodes and 1, 285 interactions, including 76 guide genes along with their first degree interactors, are extracted from the predicted network (Figure 9). Supplementary Table S6 lists unique list of 76 proteins along with their direct interactors and their respective functional annotations. Among 670 proteins, a total of 508 proteins are obtained to have their respective pathway annotation which revealed the pathways related to secondary metabolite synthesis are highly enriched. Phenylpropanoid biosynthesis pathway is the most enriched including 77 proteins followed by terpenoid backbone biosynthesis, steroid biosynthesis pathway etc. listed successively in Supplementary Table S6. Similarly, GO annotations were also mined for all the 670 proteins. A total of 1, 236 GO terms could be successfully associated with 492 proteins. Table S3 lists GO enrichment along with corresponding description of each GO term. As expected GO terms related to primary and secondary metabolites biosynthetic pathways are highly enriched. Protein Ote100115070041 is found to have highest degree, interacting with 91 other partners and is found to be enzyme Taxane 10-beta-hydroxylase an oxidoreductase involved in biosynthesis of taxol. Taxol is an antimitotic agent that prevent range of cancers like carcinoma, scrcoma and melanoma^70^. Similarly, protein Ote100174760081 having 70 direct interactors is the next highest degree node in the metabolic protein subnetwork of *O. tenuiflorum*. This protein is also found to be a farnesyl-pyrophosphate synthase that is involved in biosynthesis of linalool. Linalool possess anti-infective properties and has wide roles as an antiallergic, antiinflammatory, cleaning and perfumery agent^71^. Protein Ote100256650051 having degree 61 is predicted to be geraniol dehdrogenase that is involved in citral biosynthesis and is used as an antiseptic agent. Ote100233130031 having degree 59 is identified as farnesyl-pyrophosphate synthase followed by Ote100265920011 with 58 links and is found to be enzyme r-linool_synthase involved in biosynthesis of linalool.

**Figure 9:**
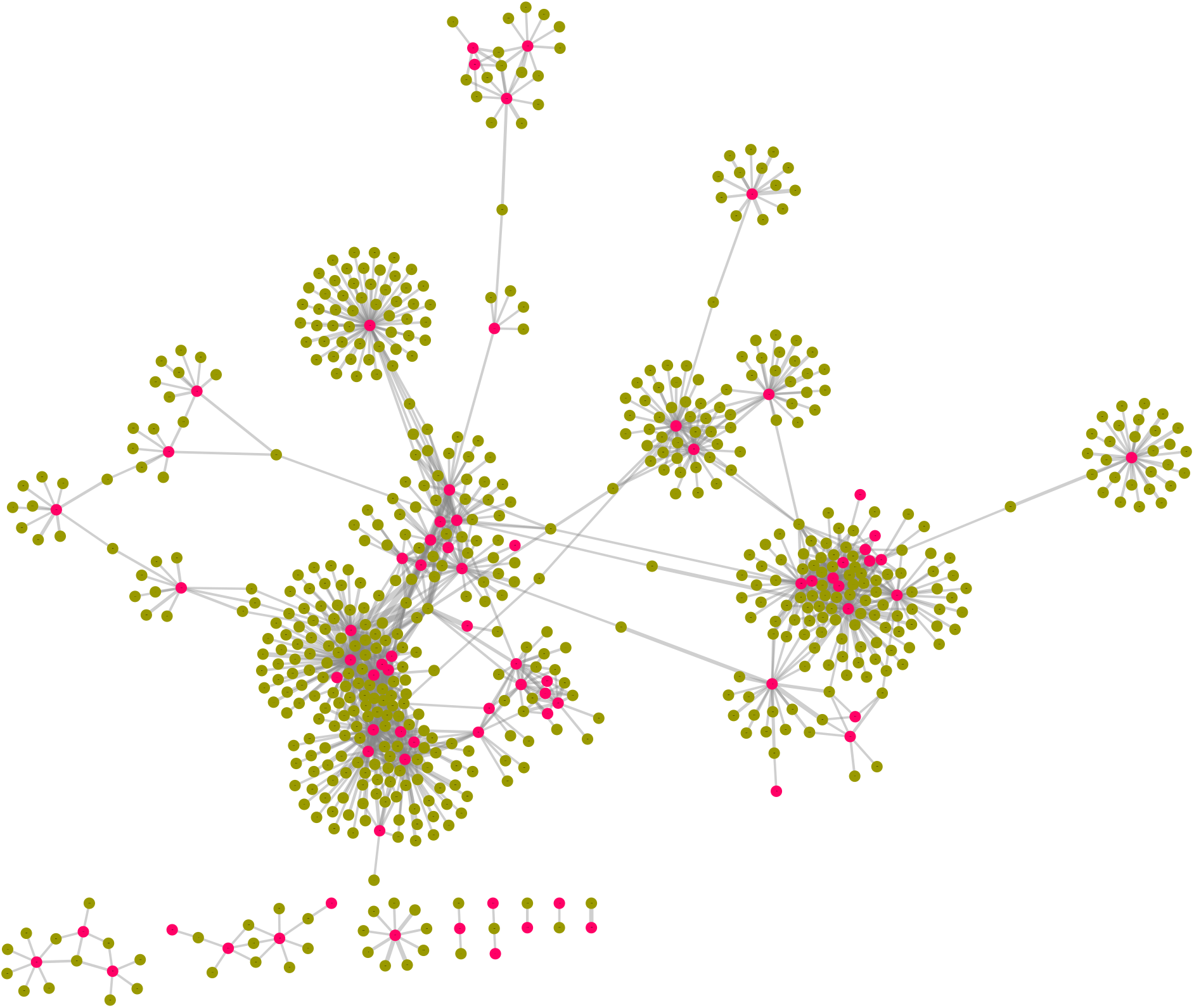
A subnetwork of 76 guide genes involved in selected 12 metabolite biosynthetic pathways of *O. tenuiflorium* comprising of 670 nodes and 1, 285 interactions. Guide genes are highlighted in pink color.

## Summary

The holy basil, tulsi, has a rich history from the ancient times of Ayurveda to be used as a cure for large number of diseases and disorders. The botanical is used extensively in a range of daily household chores like flavor enhancer of foodstuff (culinary) or being used in religious rituals. Also, it is globally used in pharmaceutical, cosmetic and perfumery industries. Based on recent genome report of tulsi, we attempted to construct and analyze its genome wide protein-protein interaction network in order to understand various regulatory mechanisms and pathways. Interactome maps allow us to explore complex genetic circuitry regulating plant responses in varying conditions and enable its sustained growth and development. In this study, we propose a robust methodology for construction and analysis of PPI networks of plants in general. Using publicly available PPI information of 49 plant species an interolog based interactome map of tulsi (TulsiPIN) consisting of 13, 660 nodes and 327, 409 interactions is developed. To the best of our knowledge this is the largest plant specific data that has been used to construct an exhaustive genome wide interologous PPI network for any plant species so far. Our methodology is essentially two fold, an interologous based PPI network is first constructed and then domain domain interactions were utilized to mark the confidence of each predicted interaction. By combining these two methods, we associate a score with each interaction in the TulsiPIN to construct a high confidence PPI network (hc-TulsiPIN) consisting of 7, 719 nodes and 95, 532 binary pairs of interactions.

Biological significance or function based reliability of each predicted interolog was assessed in terms of two measures viz. FHI and PCS that utilize GO based functional terms and subcellular localization information of each protein. Most of the interologs are found to have high values of both FHI and PCS as compared to random ensembles. We also propose a novel methodology for identification of key proteins in a PPI network by combining three architectural measures namely degree, betweenness and eigenvector centralities. A total of 1, 625 statistically significant key proteins are identified form hc-TulsoPIN. Further, functional modules are identified from hc-TulsiPIN, which revealed compliance in terms of GO based function similarity index and colocalization similarity index. A total of 15 modules are finally selected and characterized for their pathways. Most of the modules are found to contain proteins involved in similar functions, having same location and are found as intermediates of same pathways. Additionally, subnetwork of proteins involved in 10 secondary metabolite biosynthetic pathways is constructed from hc-TulsiPIN and further evaluated for its functional content. The subnetwork revealed that most of the interacting proteins are involved in terpene biosynthesis. Moreover, the modular architecture of TulsiPIN is assessed and it is found that proteins involved in a module are involved in similar functions.

As our hc-TulsiPIN provides a comprehensive map of interacting proteins of *O. tenuiflorum*, we believe, it will play an important role to understand the mechanisms underlying complex biological processes ranging from development to defense against pathological conditions in this medicinally important herb. Although, hc-TulsiPIN represents a subset of the actual network functioning *in vivo*, however, it surely will provide a rich source of information on protein functioning in tulsi to the scientific community. We believe that the proposed methodology can be applied to other plants as well for which interactomic information is still lacking.

## Acknowledgement

We thank Central University of Himachal Pradesh for providing the required infrastructure and computational facilities. VS^*†*^ thanks Council of Scientific and Industrial Research (CSIR), India for providing Junior Research Fellowship (JRF).

## Author Contributions

VS^*∗*^ conceptualized and designed the research framework as well as supervised the entire study. VS^*†*^ performed the computational experiments. VS^*†*^, GS and VS^*∗*^ analyzed the data and interpreted results. VS^*†*^ and VS^*∗*^ wrote and finalized the manuscript.

